# Exploration of deep-learning based classification with human SNP image graphs

**DOI:** 10.1101/2021.10.01.462710

**Authors:** Chao-Hsin Chen, Kuo-Fong Tung, Wen-Chang Lin

## Abstract

**Background:** With the advancement of NGS platform, large numbers of human variations and SNPs are discovered in human genomes. It is essential to utilize these massive nucleotide variations for the discovery of disease genes and human phenotypic traits. There are new challenges in utilizing such large numbers of nucleotide variants for polygenic disease studies. In recent years, deep-learning based machine learning approaches have achieved great successes in many areas, especially image classifications. In this preliminary study, we are exploring the deep convolutional neural network algorithm in genome-wide SNP images for the classification of human populations.

**Results:** We have processed the SNP information from more than 2,500 samples of 1000 genome project. Five major human races were used for classification categories. We first generated SNP image graphs of chromosome 22, which contained about one million SNPs. By using the residual network (ResNet 50) pipeline in CNN algorithm, we have successfully obtained classification models to classify the validation dataset. F1 scores of the trained CNN models are 95 to 99%, and validation with additional separate 150 samples indicates a 95.8% accuracy of the CNN model. Misclassification was often observed between the American and European categories, which could attribute to the ancestral origins. We further attempted to use SNP image graphs in reduced color representations or images generated by spiral shapes, which also provided good prediction accuracy. We then tried to use the SNP image graphs from chromosome 20, almost all CNN models failed to classify the human race category successfully, except the African samples.

**Conclusions:** We have developed a human race prediction model with deep convolutional neural network. It is feasible to use the SNP image graph for the classification of individual genomes.

## Background

Artificial intelligence or deep learning is a rapid evolving machine learning research field in recent years, especially extended applications of deep neural network based machine learning algorithm in many diverse areas [1-6]. There are more and more applications and new advancements reported in various topics utilizing deep learning models, such as image classification [7-13]. Image classification has achieved record breakthrough and advancement in just a few years since the adaptation of deep neural network into this subject. Advance of deep convolutional neural network (CNN) certainly improves prediction performance and generate new success and extended applications for computer vision and image recognition areas. Such applications have been greatly utilized in daily life with enormous impacts, such as facial recognition, smart phone systems, and automatic driving assisted technologies. In biomedical researches, deep convolutional neural network has significantly promoted in medical imaging fields and their clinical applications possess tremendous potentials in forthcoming clinical medicines [7, 8, 10]. It is anticipated that artificial intelligence-based diagnostic decision support powered by deep learning algorithm would have great influence in future medical imaging applications such as X-ray images, computed tomography (CT)/ Magnetic resonance imaging (MRI) scan images, and histopathological images [14-16]. It is likely to see more utilizations of deep learning algorithm moved into the medical genomic researches and applications [2].

In recent years, completion of human genome sequences rapidly accelerates the discovery of human disease-associated genes. Next generation sequencing (NGS) platform certainly is a remarkable technological innovation in the modern genomic researches and biomedical applications. In particular, genome-wide association study (GWAS) is another significant development in human genetic researches and has been greatly utilized in many successful studies to identify important disease associated traits [17-19]. Besides whole genome sequences, GWAS serves as an alternative cost-effective approach in the heritable disease gene interrogations and mostly relied on the single nucleotide polymorphic variants in the human genome sequences. These single nucleotide polymorphism (SNP) variants occur around one SNP per thousand bases throughout the human genome. Currently, there are several hundred millions of SNPs reported. With the wide application of NGS technology, we are anticipating more SNPs or personal variants being sequenced and discovered. In addition, increased numbers of sequenced human individual genome are accumulated through the efforts of BioBanks and other large-scale sequencing projects [20-23]. Therefore, it is possible to use very large numbers of human population for the genome-wide polygenic score studies of complex human traits [24-26]. Recently, there are significant findings in using such polygenic analysis approaches for the prediction of human heights and other complex phenotypes [27]. It would be beneficial to explore the applications of deep learning neural network methods in such a challenging area of complex genetic traits with very large numbers of genomic sequence variants.

While deep learning-based neural network type machine learning approaches have achieved great success in last few years in many different fields, the utilization of convolutional neural network machine learning methodology in human genetic studies has been limited thus far [28]. In particular, DNA sequence features and closely associated biological properties using the convolutional neural network algorithm were interrogated previously [29-32]. There are reports on the whole genome scale applications due to the large genome sizes and complexities involved [25, 33]. There are millions of variants (SNPs) existed in the genome sequences of each person. In most analysis, usually only the major alleles were considered in order to reduce computation loads. Thus, minor allelic changes or personal mutations would be filtered or omitted during the analysis. Herein, we have used the SNP information from the 1000 genomes project [34], which provides high-quality SNP information of 2,504 individuals from different human populations worldwide. In this preliminary study, we would like to explore the examination of complete SNP variation information in one chromosome by using deep learning based image classification models.

## Results

### Chromosome 22 image graph

We are interested to use the deep CNN algorithm and SNP images for human genome classification. We have obtained the 1000 genome project SNP variants dataset. The 1000 genome project is aimed to create the largest public catalog of human variation and genotype data [34]. It contained genome variation sequence information of more than 2,500 individuals. They came from various ethnic backgrounds (26 populations of five major ethnic groups) (Table 1). In total, there are more than 84 million SNPs, 3.6 million insertions/deletions (indels), approximately 60 thousand structural variants, and associated genotypes collected already. This large amount of data can improve and address the studies of human genome complex diversities. Due to the large numbers of SNPs and computation load consideration, we did not use all 84 million SNPs for the image graph generation. In this initial feasibility study, we selected chromosome 22 as the test dataset for generating image graphs because of the small size of chromosome 22. There are around one million SNP sites reported from the 1000 genome project (1,060,388 SNPs).

**Table 1.**
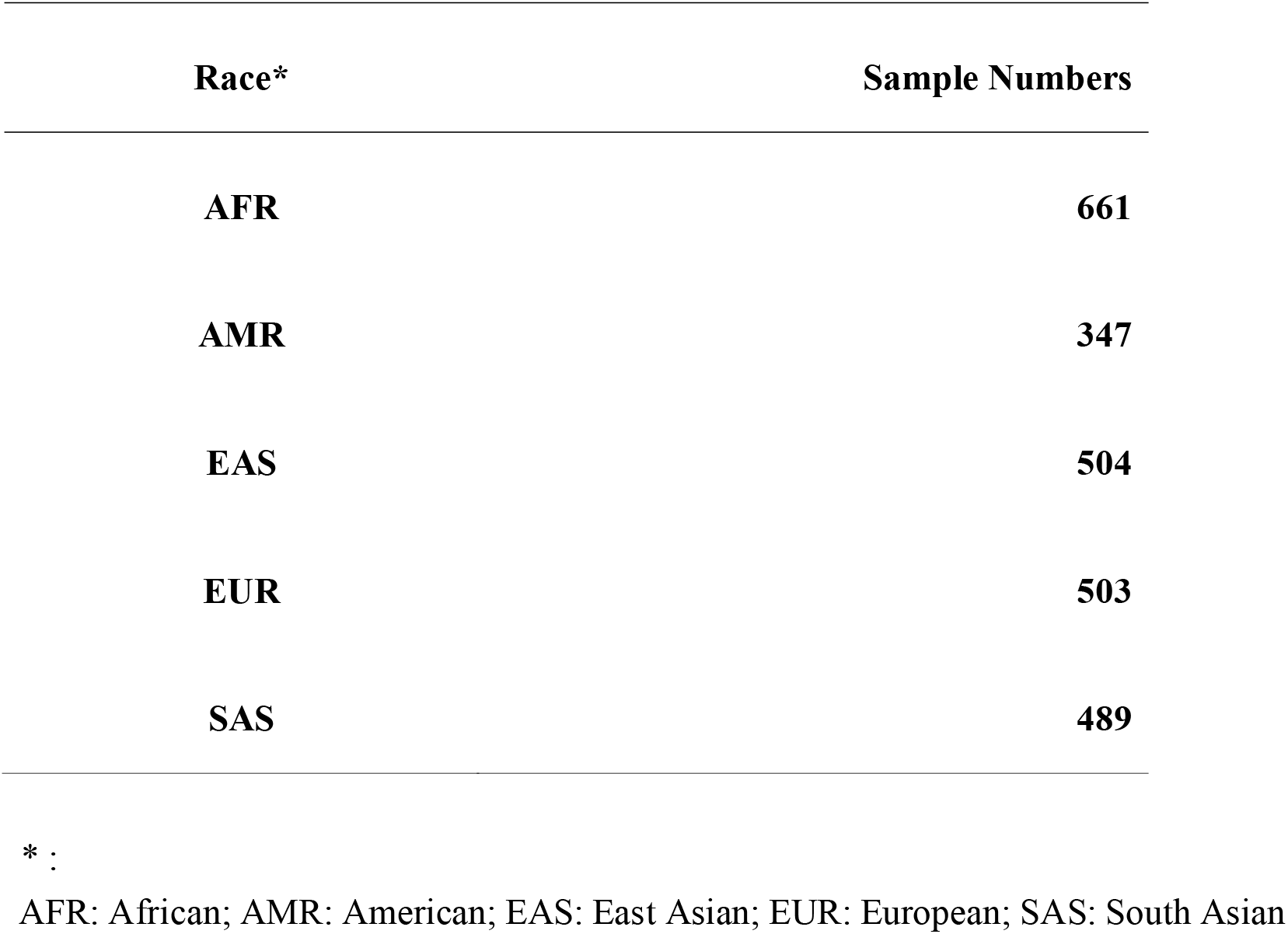
1000 genome project dataset. Five major human races listed in the 1000 genome projects and the numbers of samples.

| **Race\*** | **Sample Numbers** |
| --- | --- |
| **AFR** | **661** |
| **AMR** | **347** |
| **EAS** | **504** |
| **EUR** | **503** |
| **SAS** | **489** |

\* :
AFR: African; AMR: American; EAS: East Asian; EUR: European; SAS: South Asian

We then generated a SNP image graph from the 1000 genome project data. All samples were converted to such SNP image graphs by a customized python processing program. A typical SNP image graph is illustrated in Figure 1, which is around 1030 × 1030 pixels and 165 kilobytes in file size. Inside the graph, each pixel matches to a specific SNP type at a corresponding chromosome position. In the initial experiment, we generated the SNP image graph line by line to form a two-dimensional square image. Each line will contain 1030 SNPs, and subsequent SNP will continue to draw in the next line. We also used 16 different colors to represent the different SNP types: AA-red; AT-purple; AG-yellow; AC-black; TA-orange; TT-purple; TG-pink; TC-dark green; GA-dark salmon; GT-deep lilac; GG-green; GC-cyan; CA-sky blue; CT-summer sky; CG-dark green; CC-blue. There is no specific graphic pattern observed in the first glance between five races (Figure 1 and Supplementary Figure 1).

**Figure 1.**
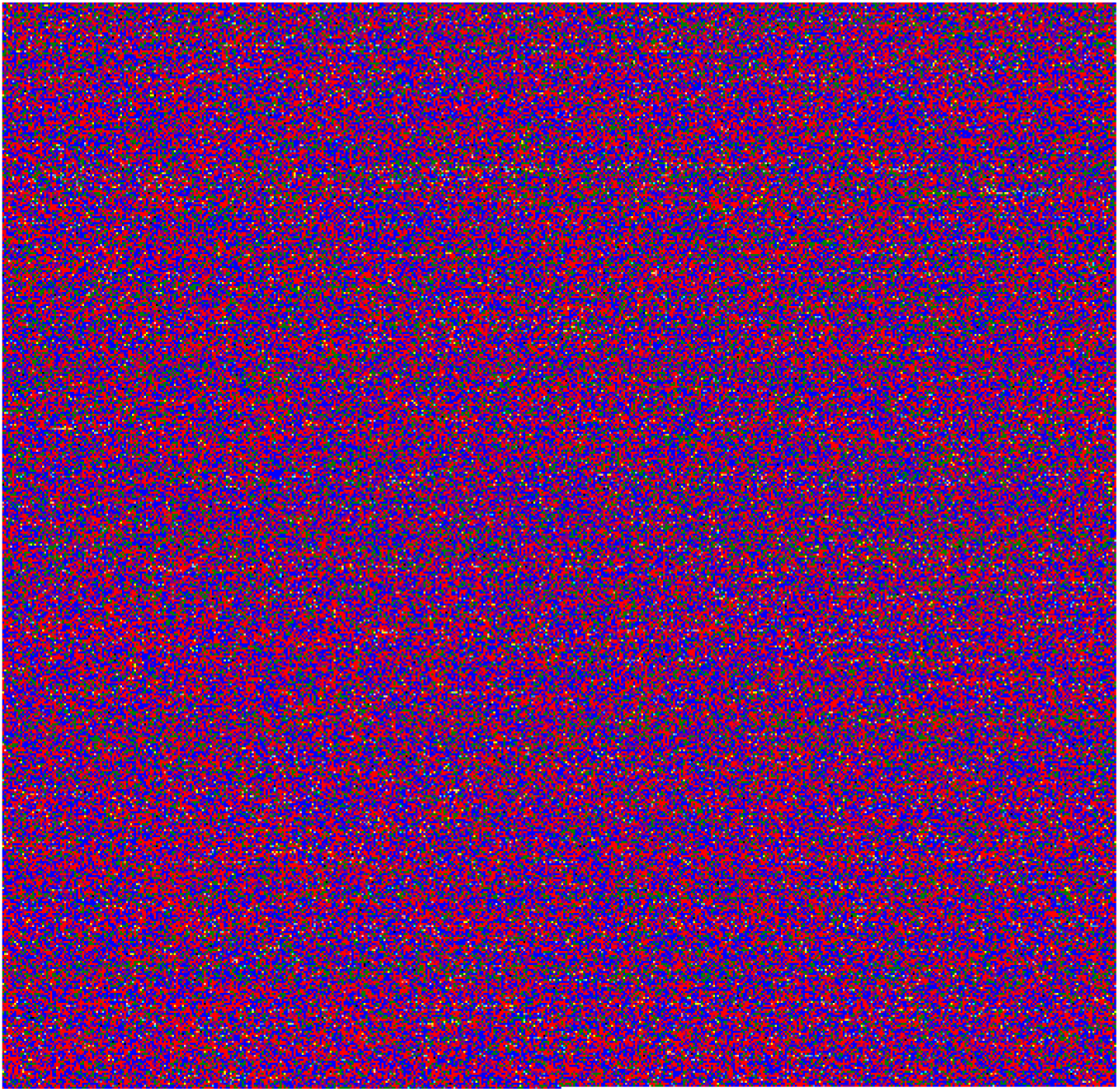
A representative SNP image graph of chromosome 22. The chromosome 22 SNP data from AFR HG01879 was used. 1,060,388 SNP records of chromosome 22 were converted into a 1030 × 1030 color image graph. Each pixel matches to a specific SNP genotype type at a corresponding chromosome position. This simple two-dimension graph was generated with line by line drawing of each SNP genotypes and 16 different colors were used as indicated in the Methods section.

### Classification of human race by deep learning algorithm

The deep convolutional neural network has achieved excellent results in the image classification area. Recently, it has been applied in the biomedical image applications with great potential in future clinical medicine practices, such as CT and x-ray images. In this experimental study, we attempted to use the deep convolutional neural network for classifying human SNP images. Among the five major human races, we first randomly selected and reserved the SNP image graphs of 30 individual for the validation phase and used the rest of image graphs for the training phase of deep learning models. There is some difference between the numbers of SNP image graphs available for each human races (Table 1). Thus, for the training dataset (total of 2,354 individuals), the numbers of image graphs used are 631 graphs for AFR; 317 graphs for AMR; 474 graphs for EAS; 473 graphs for EUR; 459 graphs for SAS.

There are several popular deep CNN algorithms available, we have tested VGG16 and ResNet 50. ResNet 50 seems to have good performance and acceptable computation time in our preliminary test run. We have repeated the deep learning experiment three times with 10 folds cross-validation, where 10% of testing graphs were used for evaluation after the deep learning model training with 90% of graphs. The final training results are very good and confusion matrix data is shown in Figures 2. The F1 scores are 95%, 98% and 99% for the three different independent tests (CV1 to CV3). We then used the reserved 150 SNP image graphs (30 randomly selected graphs for five races) for validating the trained CNN models. The overall accuracy is 95.8% for the independent validation test (Table 2). High accuracy was obtained with the AFR, EAS and SAS populations. Interestingly, misclassification was observed between the AMR and EUR populations. From this study, we often see the classification of AMR samples as EUR population or vice versa. It is possible that this phenomenon could result from the South Americans with European descent in the 1000 genome project sample collection.

**Table 2.**
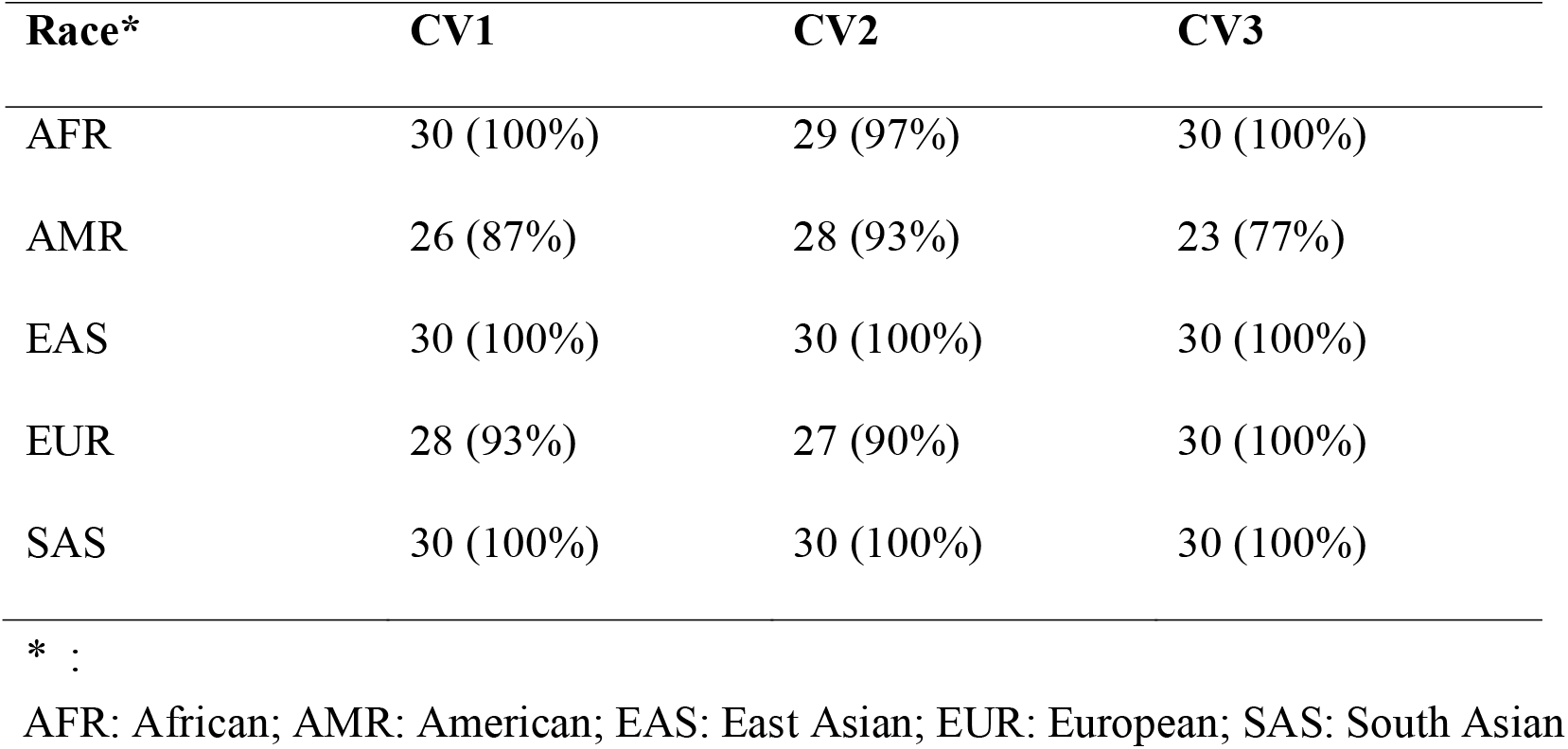
Validation Results of deep CNN models trained with 16 Category SNP images. CV1, CV2 and CV3 are three independent tests.

| **Race\*** | **CV1** | **CV2** | **CV3** |
| --- | --- | --- | --- |
| AFR | 30 (100%) | 29 (97%) | 30 (100%) |
| AMR | 26 (87%) | 28 (93%) | 23 (77%) |
| EAS | 30 (100%) | 30 (100%) | 30 (100%) |
| EUR | 28 (93%) | 27 (90%) | 30 (100%) |
| SAS | 30 (100%) | 30 (100%) | 30 (100%) |

\* :
AFR: African; AMR: American; EAS: East Asian; EUR: European; SAS: South Asian

**Figure 2.**
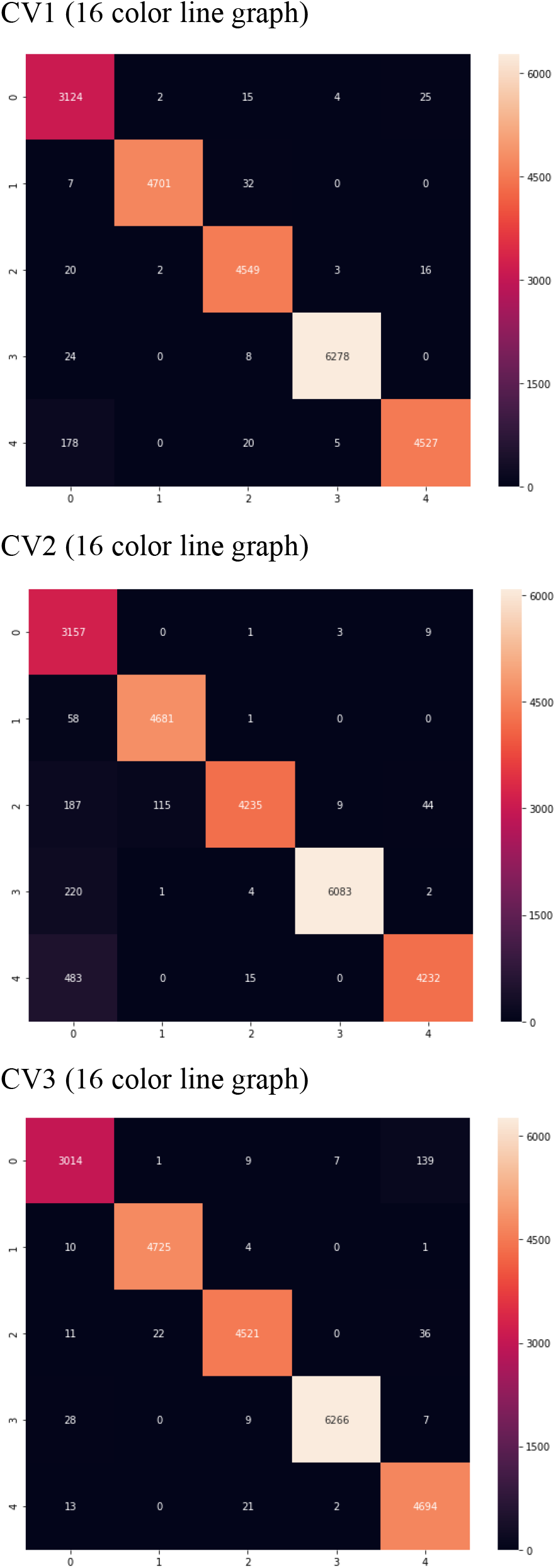
Confusion matrix plots produced by the trained deep CNN classifier with 16 color line by line SNP image graphs. The five category labels are the human races used in the study: 0-AMR: American; 1-EAS: East Asian; 2-SAS: South Asian; 3-AFR: African; 4- EUR: European. True label categories are in the Y-axis and predicted label categories are in the X-axis. The numbers in each element indicated the positive matches of predicted graphs. CV1 (A), CV2 (B) and CV3 (C) represents the results from three independent tests.

### Exploration with reduced SNP color levels and spiral images

In the preliminary examination, we used 16 colors to represent the biallelic SNP composition of human diploid genomes. However, most SNP information is not allele-specific due to the sequencing library preparation, therefore, we also inspect reduced color image levels to represent the SNP image graph. In this investigation, 8 different colors were used: AA/TT-blue; AT/TA-black; AG/GA-green; AC/CA-white; TG/GT-yellow; TC/CT-brown; GG/CC-red; GC/CG-orange. The representative SNP image graph is shown in Supplementary Figure 2. We then applied a comparable deep learning pipeline to classify all 1000 genome project samples using the 8-color image graphs. It is interesting to note that the training models still maintain good accuracy. The F1 scores are 89%, 83% and 88% in three different tests (Supplementary Figure 3). Again, we used 150 reserved randomly selected SNP image graphs for the independent validation. As shown in Supplementary Table 1, the overall accuracy is 91.1%. Misclassification was observed mainly in the AMR population.

In previous tests, we draw the SNP images using line by line scheme according to the chromosomal position of SNPs. To interrogate the chromosome positional effects, we then tested the SNP image graphs generated by a spiral pattern. We generated the spiral SNP image graphs and then trained new models. In three independent tests, the F1 scores were 98%, 97% and 98% (Figure 3). Validation test was performed again with the 150 reserved SNP image graphs. Comparable accuracy was obtained, the overall accuracy is 96% (supplementary Table 2). High accuracy was found in the AFR, EAS and SAS populations. Misclassification was observed again between the AMR and EUR populations. This suggests that overt SNP image graphs generated by line or spiral images might be essential for the deep CNN models.

**Figure 3.**
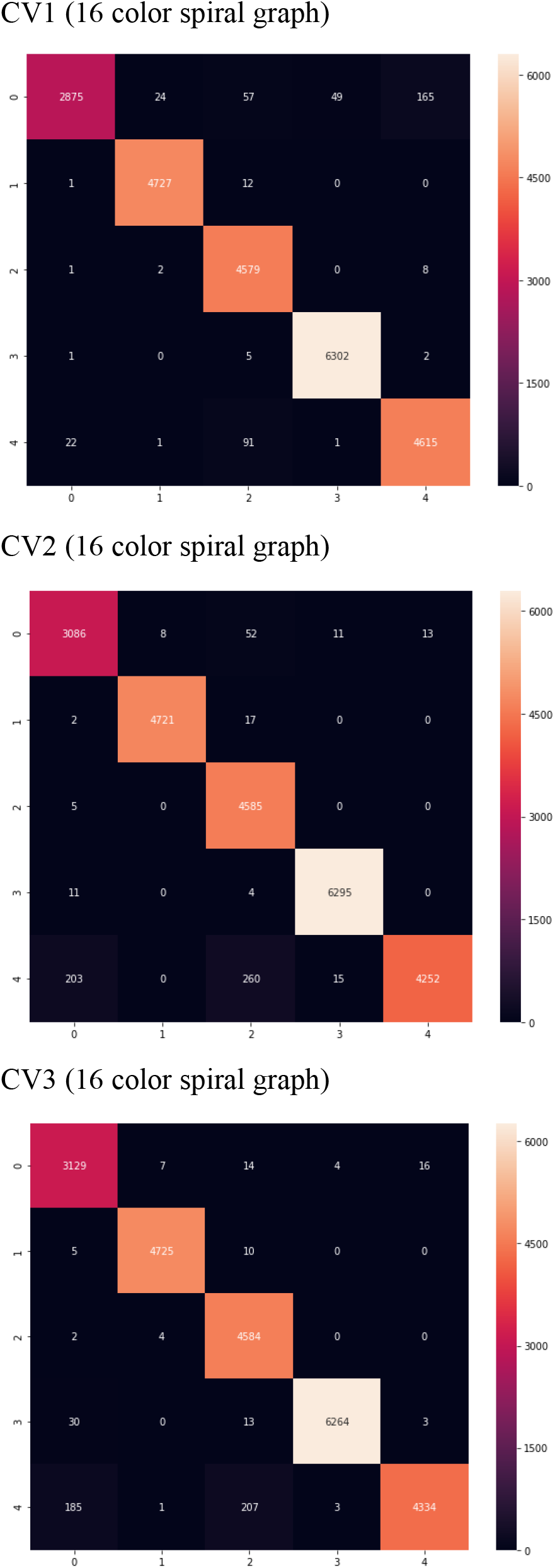
Confusion matrix plots produced by the trained deep CNN classifier with 16 color spiral SNP image graphs. The five category labels are the human races used in the study: 0- AMR: American; 1- EAS: East Asian; 2- SAS: South Asian; 3- AFR: African; 4- EUR: European. True label categories are in the Y-axis and predicted label categories are in the X-axis. The numbers in each element indicated the positive matches of predicted graphs. CV1, CV2 and CV3 represents the results from three independent tests.

### Chromosome 20 evaluation with the trained models from chromosome 22

Finally, we tested the CNN models trained from chromosome 22 for the classification with SNP image graphs from other chromosomes. We selected chromosome 20 as the initial test data. As expected, the validation results failed and almost no samples predicted correctly except for the AFR population. Intriguingly, all AFR samples were still predicted correctly (data not shown). It is possible that there is a SNP variation (image patterns) preserved in the ancestral AFR population in the human lineage, which would require further investigations.

## Discussion

The 1000 genome project is the prime resource of human population studies with high quality genome sequence variation information in diverse ethnic backgrounds. Using the chromosome 22 SNP information of more than 2,500 individuals, we have developed a human race prediction model by a deep convolutional neural network method. It performed well and showed 95% classification accuracy with the five major human race categories. This demonstrated that it is possible to use the current highly advanced deep convolution neural network models in classifying large numbers of human genomic SNP information. We then tried to evaluate the subpopulation classification of 26 human races listed in the 1000 genome project. The prediction results dropped considerably with the F1 score of 75% (Supplementary Figure 4). This could due to the small sample sizes in the training dataset from 1000 genome project in the 26 population category. This demonstrates the critical issue of training sample volume. However, with more and more genomic sequences available, it is not a crucial consideration. On the contrary, it is the reason we performed this preliminary study due to the large numbers of human genome sequences released in many BioBank resources and integrated international disease consortiums. Our efforts here suggest that it is feasible to use the popular deep learning convolution neural network models in very large scale human SNP genotypes.

One significant observation here is the misclassification between the AMR and EUR populations. It is possible that this indicates the South Americans with European descent in the 1000 genome project sample collection. In a recent report [35], excess European ancestry SNP signatures were found in analyzed Latin America populations (Colombia, Mexico, Peru, and Puerto Rico). Puerto Rico and Colombia also have the highest levels of European ancestry (72% and 64%), while Mexico has slightly less European ancestry (48%). Peru samples showed only 18% of European ancestry. This report in some way supports our speculation. Of course, we have not evaluated that this result is only unique to the chromosome 22 SNP image graphs. It is possible that we could have better classification results with the inclusion of other chromosome SNP information. The 1000 genome project provides 84 million SNPs and could generate a large SNP image graph of 9200 pixels X 9200 pixels image, which could have more specific pattern signature information.

Utilization of large numbers SNPs in deep learning based genomic analysis is just launched, more studies and explorations are needed. An accurate human height predictor was reported by analyzing over 630,000 SNPs of close to 500,000 individuals, which would comprise over 20k informative SNPs in the predictor [27]. In a recent 2018 report by Bellot et al [36], comparison of Convolutional Neural Network (CNNs) and Multilayer Perceptrons (MLPs) machine learning approaches in UK BioBank analysis did not find any significant advantages of these deep learning methods over the Bayesian linear regression model. However, only less than 50,000 SNPs were used in this report and the CNN model used had only three connected layers and one convolutional layer. Our study here provides a new research area for the applications of deep learning based image classifiers in human genomic sequences. Since there are many successful deep learning algorithms developed for the image analysis already, by converting the SNPs information into a global SNP image graph, we could greatly improve the human genome classification and complex trait discovery processes.

## Conclusion

We have developed a human race prediction model based on the ResNet deep convolutional neural network model and one million chromosome 22 SNP information. In the future, it is feasible to use the SNP image graph for the major human race classification of individual samples.

## Methods

### Samples collection from 1000 genome project

We have obtained phase 3 SNP information from the 1000 genome project [34]. Due to the purpose here is to exploratory how deep learning techniques use on SNP images, therefore, we selected chromosome 22 as the target dataset in preliminary analysis. The obtained dataset includes five major races and 26 subpopulation groups. Five races were consisted of 661 samples for African (AFR); 347 samples for American (AMR); 504 samples for East Asian (EAS); 503 samples for European (EUR); and 489 samples for South Asian (SAS). There are 26 subpopulations groups in the dataset which includes: African-Caribbean (ACB, 96 samples);African-American SW(ASW, 61 samples; Esan (ESN, 99 samples);Gambian Mandinka (GWD, 113 samples);Luhya (LWK, 99samples); Mende (MSL, 85 samples);Yoruba (YRI, 108 samples);Colombian (CLM, 94 samples);Mexican-American (MXL, 64 samples);Peruvian (PEL, 85 samples);Puerto Rican (PUR, 104 samples);Dai Chinese (CDX, 93 samples);Han Chinese (CHB, 103 samples);Southern Han Chinese (CHS, 105 samples);Japanese (JPT, 104 samples);Kinh Vietnamese (KHV, 99 samples);Bengali (BEB, 86 samples);Gujarati(GIH, 103 samples);Indian (ITU, 102 samples); Punjabi (PJL, 96 samples);Sri Lankan (STU, 102 samples);CEPH(CEU, 99 samples);Finnish(FIN, 99 samples);British(GBR, 91 samples);Spanish(IBS, 107 samples);Tuscan(TSI, 107 samples).

### SNP information and filters

Using bcftools tool which suggested by 1000 genomes project, we extracted chromosome 22 variant information (2,504 samples with 1,103,547 SNP IDs). We filter the variant genotype data to retain only the single nucleotide polymeric variants and remove other variations (Indels and other structure variants). Total of 1,060,388 SNP records was used for subsequent analysis. For chromosome 20, 1,812,841 SNP IDs obtained in total and we selected 1,746,157 SNP records for test data with the same filter criteria.

### SNP image graph generation

We used 1,060,388 SNP records to generate two types of a square matrix graphs (sequentially aligned by either line or spiral format) in order to build simple two-dimension graphs for each human genome sample. Additionally, we used zero to padding insufficient records for creating 1030 × 1030 square matrix. We then first used 16 different colors to indicate each bi-allelic genotype information from the 1000 genome project. We used 16 colors to represent the different SNP types: AA-red; AT-purple; AG-yellow; AC-black; TA-orange; TT-purple; TG-pink; TC-dark green; GA-dark salmon; GT-deep lilac; GG-green; GC-cyan; CA-sky blue; CT-summer sky; CG-dark green; CC-blue. In further analysis, due to the specific allelic variants are not specified in 1000 genomes project dataset, we also tried to use 8 colors for reducing image complexity levels in generating SNP image graphs. The eight colors used were AA/TT-blue; AT/TA-black; AG/GA-green; AC/CA-white; TG/GT-yellow; TC/CT-brown; GG/CC-red; GC/CG-orange. Based on the 1000 genome project dataset, we categorized generated image graphs into either 5 major races or 26 subpopulation groups in subsequent training for deep CNN models.

### Deep CNN model training

The deep learning model training was performed on a GPU (Nvidia GeForce RTX 2080) workstation with two Intel Xeon E5-2620 CPU and 128 RAM. ResNet50 algorithm was employed for deep convolutional neural network model training in this study. The typical training phase is around 48 hours. ResNet50 has 50 layers and we added regularizer and dropout layers for eliminating overfitting (learning rate=e-4; dropout=0.4; epochs=20 and batch_size=32). We randomly selected 30 sample graphs in each race for further model validation phase. For the rest of image graphs, 10% of graphs in each race were used as a testing set for evaluating 90% of the graphs for model training. In the model training process, we randomly cropped each image graph 100 times into 256 × 256 pixels for fitting the ResNet model, and then repeated the model training experiment 3 times with this 10-fold cross-validation. Three independently trained CNN models were tested (CV1, CV2 ad CV3). We used ‘relu’ activation and adjusting the learning rate and epochs for the best fit of model training. Total parameters in the neural network training phases were more than 25,000,000. Regards the evaluation metrics, accuracy and F1 score were employed to this study. Accuracy is an overall evaluation of how accurate the categories were predicted, which is calculated by the numbers of correct prediction over the total number of predictions. F1 score can gain the weighted average of the precision and recall to interpret the result more objectively. We also performed the confusion matrix analysis to obtain a better understanding of the complete performance of the model.

In the 26 subpopulation experiments, due to the small sample size, we have extracted 10 graphs randomly which has the most numbers of the subpopulation sample groups from each race for later model validation test. Besides, all other image processing/cropping and CNN model training were performed using the same procedure as mentioned above.

### Model validation with 150 reserved separated SNP image graphs

The reserved separate image graph (30 graphs in each major human race, 150 samples in total) were randomly cropped 50 times into 256 × 256 pixels for trained CNN model validation. We then can calculate and summarized the results of scores to each category for cropped images. Higher score showed the higher prediction possibility of respective category. Finally, we summarized the highest numbers of predicted category for each cropped image and concluded the result of human race category prediction with the major results of all 50 cropped images in each sample. Three independently trained CNN models were tested (CV1, CV2 ad CV3).

## Abbreviations

CNN: convolutional neural network
GWAS: genome-wide association study
SNP: single nucleotide polymorphism
NGS: next generation sequencing
AFR: African
AMR: American
EAS: East Asian
EUR: European
SAS: South Asian

## Acknowledgement

We are grateful to Dr. Sheng-Wei Chen at the Institute of Information Sciences, Academia Sinica for his assistance in deep learning pipelines. We also thank the generous advice and informative discussion from Dr. Chien-Hsiun Chen of the Institute of Biomedical Sciences.

## Contributions

CHC collected and processed the 1000 genome project dataset. WCL conceived and designed the experiments. CHC and KFT established the machine learning pipeline and implemented the experiments. CHC and WCL analyzed the results. CHC and WCL wrote the paper. All authors read and approved the final manuscript.

## Data Availability

The 1000 genome SNP data used to support the findings of this study were supplied by 1000 genome project and was available through its web site. The generated SNP image graph pictures are available from the corresponding author upon request.

## Figure Legends

**Supplementary Figure 1**. Representative SNP image graph of chromosome 22 from four human races. The chromosome 22 SNP image graphs (16 color) from AMR HG00551, EAS HG00403, EUR HG00096 and SAS HG01583 of the 1000 genome project are illustrated.

**Supplementary Figure 2**. An 8-color SNP image graph of chromosome 22. 1,060,388 SNP records of chromosome 22 were converted into a 1030 × 1030 color image graph. This simple two-dimension graph was generated with line by line drawing of each SNP genotypes and 8 different colors were used in this SNP image graph.

**Supplementary Figure 3**. Confusion matrix plots produced by the trained deep CNN classifier with 8 color line by line SNP image graphs. The five category labels are the human races used in the study: 0-AMR: American; 1-EAS: East Asian; 2-SAS: South Asian; 3-AFR: African; 4-EUR: European. True label categories are in the Y-axis and predicted label categories are in the X-axis. The numbers in each element indicated the positive matches of predicted graphs. CV1, CV2 and CV3 represents the results from three independent tests.

**Supplementary Figure 4**. Confusion matrix plots produced by the trained deep CNN classifier with 16 color line by line SNP image graphs. The 26 category labels are the human races used in the study: Category: ACB-0; ASW-1; BEB-2; CDX-3; CEU-4; CHB-5; CHS-6; CLM-7; ESN-8; FIN-9; GBR-10; GIH-11; GWD-12; IBS-13; ITU-14; JPT-15; KHV-16; LWK-17; MSL-18; MXL-19; PEL-20; PJL-21; PUR-22; STU-23; TSI-24; YRI-25. True label categories are in the Y-axis and predicted label categories are in the X-axis. The numbers in each element indicated the positive matches of predicted graphs.

**Supplementary Table 1.**
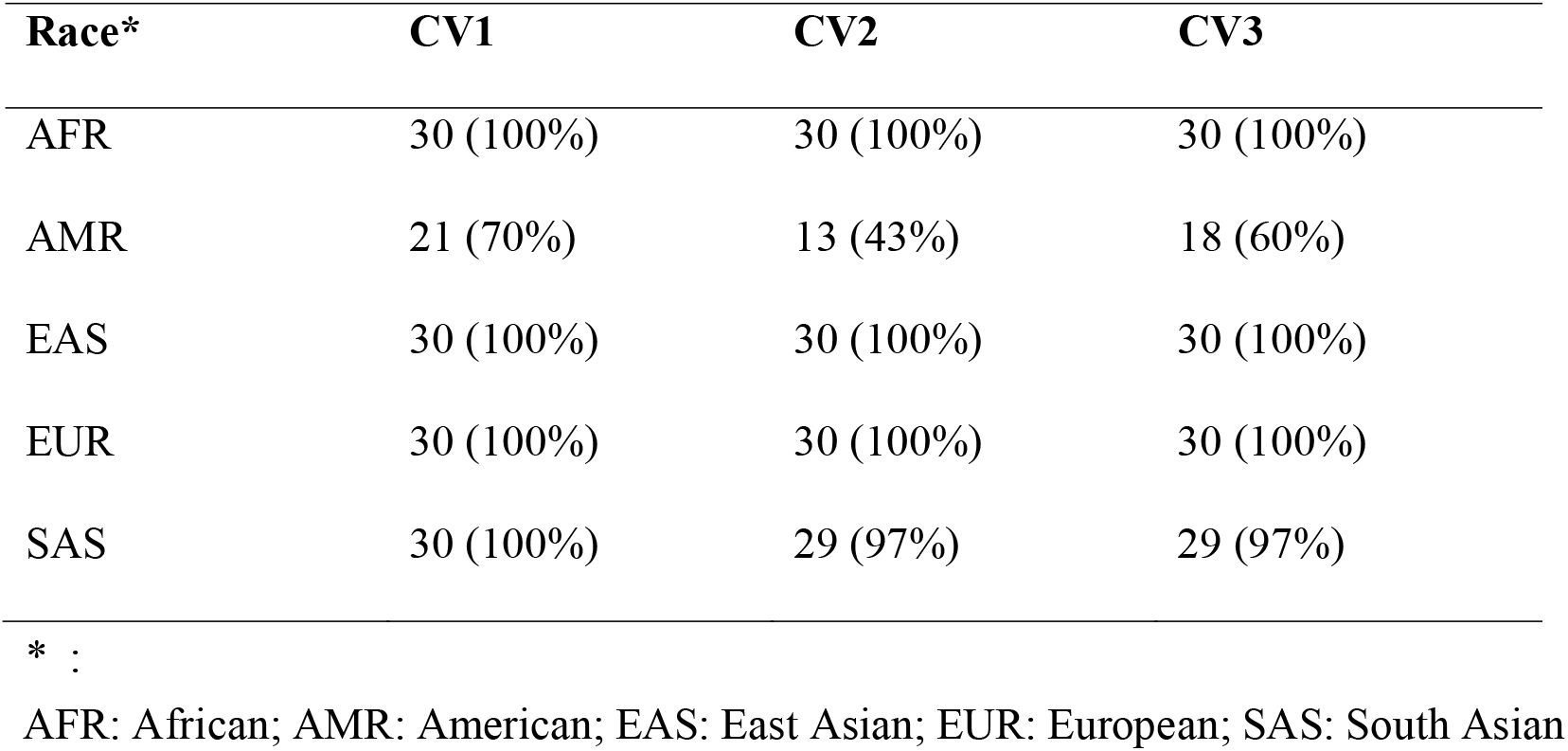
Validation Results of deep CNN models trained with 8 Category SNP images. CV1, CV2 and CV3 are three independent tests.

**Supplementary Table 2.**
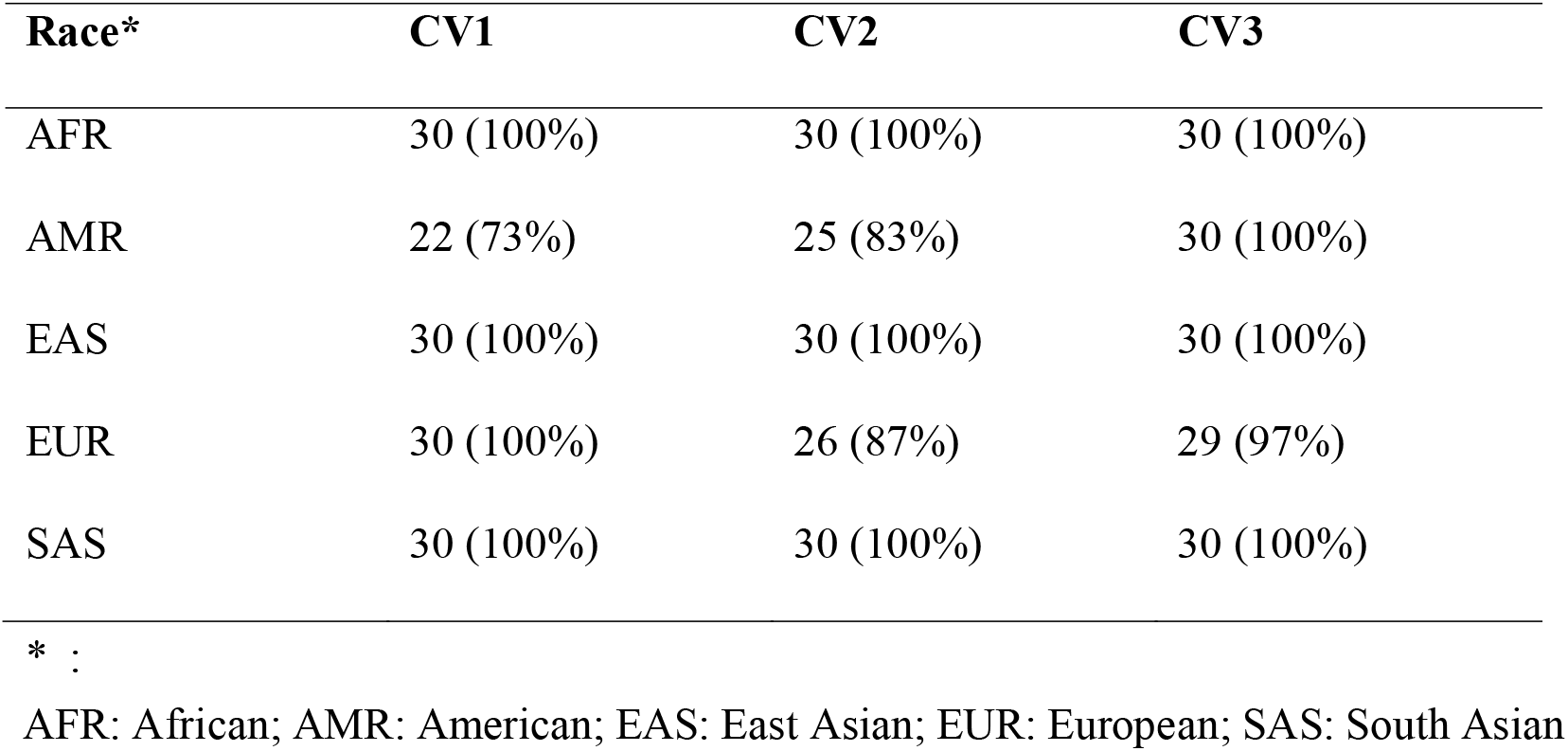
Validation Results of deep CNN models trained with spiral images. CV1, CV2 and CV3 are three independent tests.

