## Supplemental Tables and Figures for "Exploration of deep-learning based classification with human SNP image graphs"

**Supplementary Table 1. Validation Results of deep CNN models trained with 8 Category SNP images. CV1, CV2 and CV3 are three independent tests.**

| Race* | CV1 | CV2 | CV3 |
|---|---|---|---|
| AFR | 30 (100%) | 30 (100%) | 30 (100%) |
| AMR | 21 (70%) | 13 (43%) | 18 (60%) |
| EAS | 30 (100%) | 30 (100%) | 30 (100%) |
| EUR | 30 (100%) | 30 (100%) | 30 (100%) |
| SAS | 30 (100%) | 29 (97%) | 29 (97%) |

* :

AFR: African; AMR: American; EAS: East Asian; EUR: European; SAS: South Asian

**Supplementary Table 2. Validation Results of deep CNN models trained with spiral images. CV1, CV2 and CV3 are three independent tests.**

| Race* | CV1 | CV2 | CV3 |
|---|---|---|---|
| AFR | 30 (100%) | 30 (100%) | 30 (100%) |
| AMR | 22 (73%) | 25 (83%) | 30 (100%) |
| EAS | 30 (100%) | 30 (100%) | 30 (100%) |
| EUR | 30 (100%) | 26 (87%) | 29 (97%) |
| SAS | 30 (100%) | 30 (100%) | 30 (100%) |

* :

AFR: African; AMR: American; EAS: East Asian; EUR: European; SAS: South Asian

**Supplementary Figure 1.** Representative SNP image graph of chromosome 22 from four human races. The chromosome 22 SNP image graphs (16 color) from AMR HG00551, EAS HG00403, EUR HG00096 and SAS HG01583 of the 1000 genome project are illustrated.

**Supplementary Figure 2.** An 8-color SNP image graph of chromosome 22. 1,060,388 SNP records of chromosome 22 were converted into a 1030 x 1030 color image graph. This simple two-dimension graph was generated with line by line drawing of each SNP genotypes and 8 different colors were used in this SNP image graph.

**Supplementary Figure 3.** Confusion matrix plots produced by the trained deep CNN classifier with 8 color line by line SNP image graphs. The five category labels are the human races used in the study: 0- AMR: American; 1- EAS: East Asian; 2- SAS: South Asian; 3- AFR: African; 4- EUR: European. True label categories are in the Y-axis and predicted label categories are in the X-axis. The numbers in each element indicated the positive matches of predicted graphs. CV1, CV2 and CV3 represents the results from three independent tests.

**Supplementary Figure 4.** Confusion matrix plots produced by the trained deep CNN classifier with 16 color line by line SNP image graphs. The 26 category labels are the human races used in the study: Category: ACB-0; ASW-1; BEB-2; CDX-3; CEU-4; CHB-5; CHS-6; CLM-7; ESN-8; FIN-9; GBR-10; GIH-11; GWD-12; IBS-13; ITU-14; JPT-15; KHV-16; LWK-17; MSL-18; MXL-19; PEL-20; PJL-21; PUR-22; STU-23; TSI-24; YRI-25. True label categories are in the Y-axis and predicted label categories are in the X-axis. The numbers in each element indicated the positive matches of predicted graphs.

**Supplementary Figure 1.**

AMR HG00551

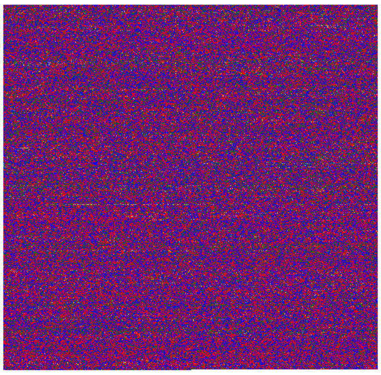

EAS HG00403

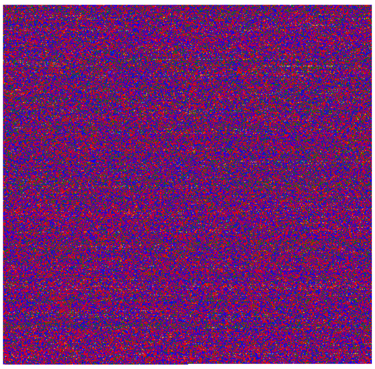

EUR HG00096

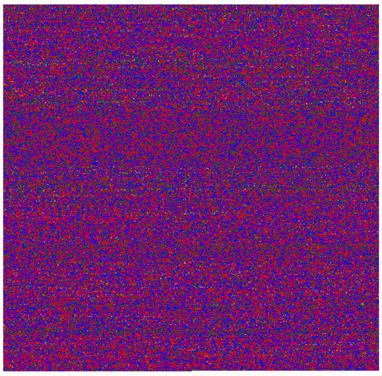

SAS HG01583

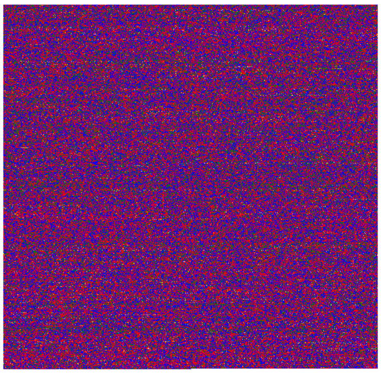

**Supplementary Figure 2.**

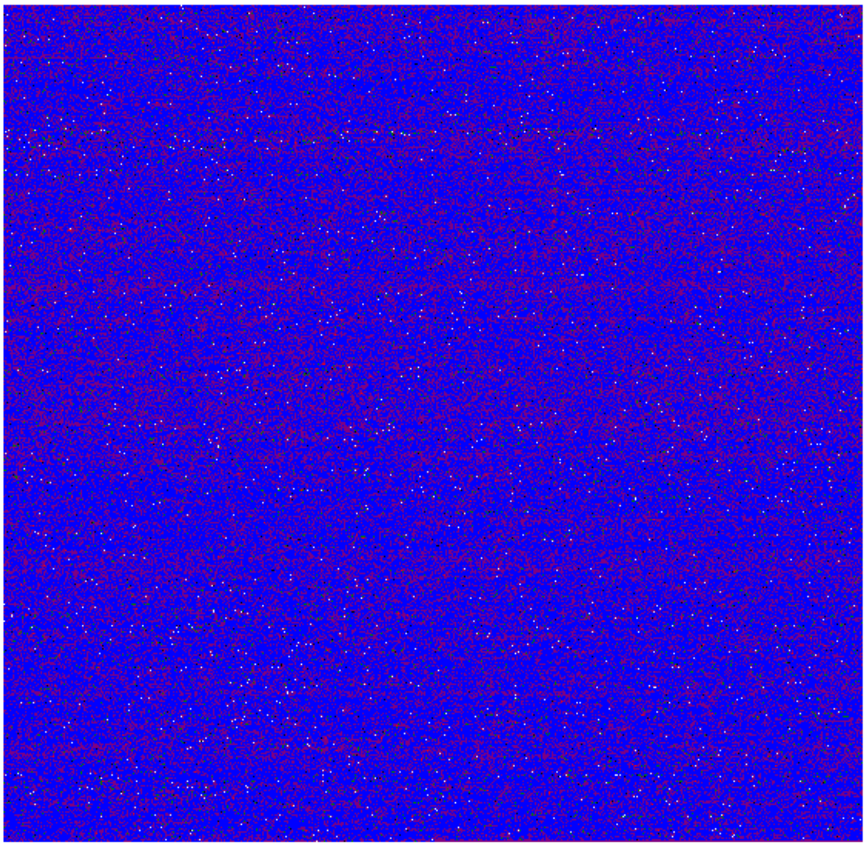

**Supplementary Figure 3**

CV1 (8 color line graph)

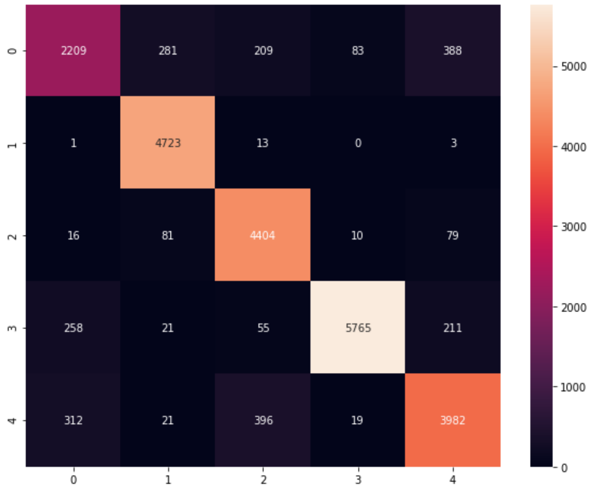


CV2 (8 color line graph)

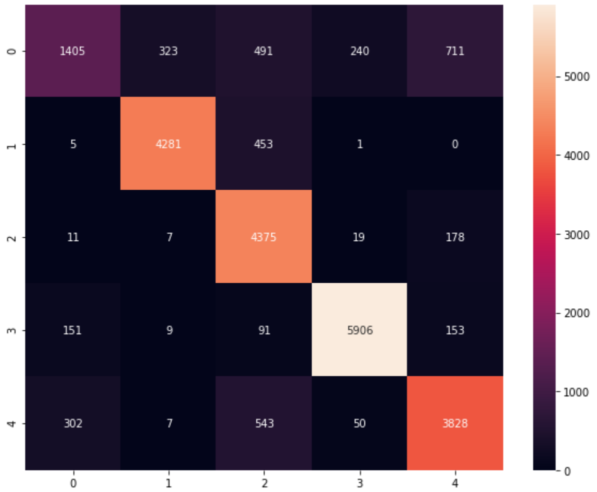


CV3 (8 color line graph)

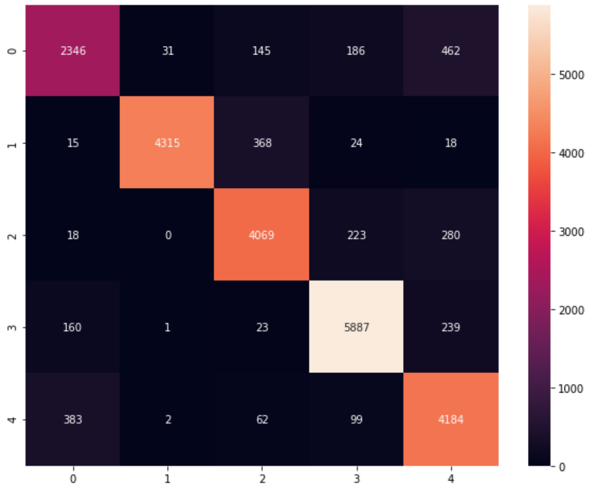

**Supplementary Figure 4**

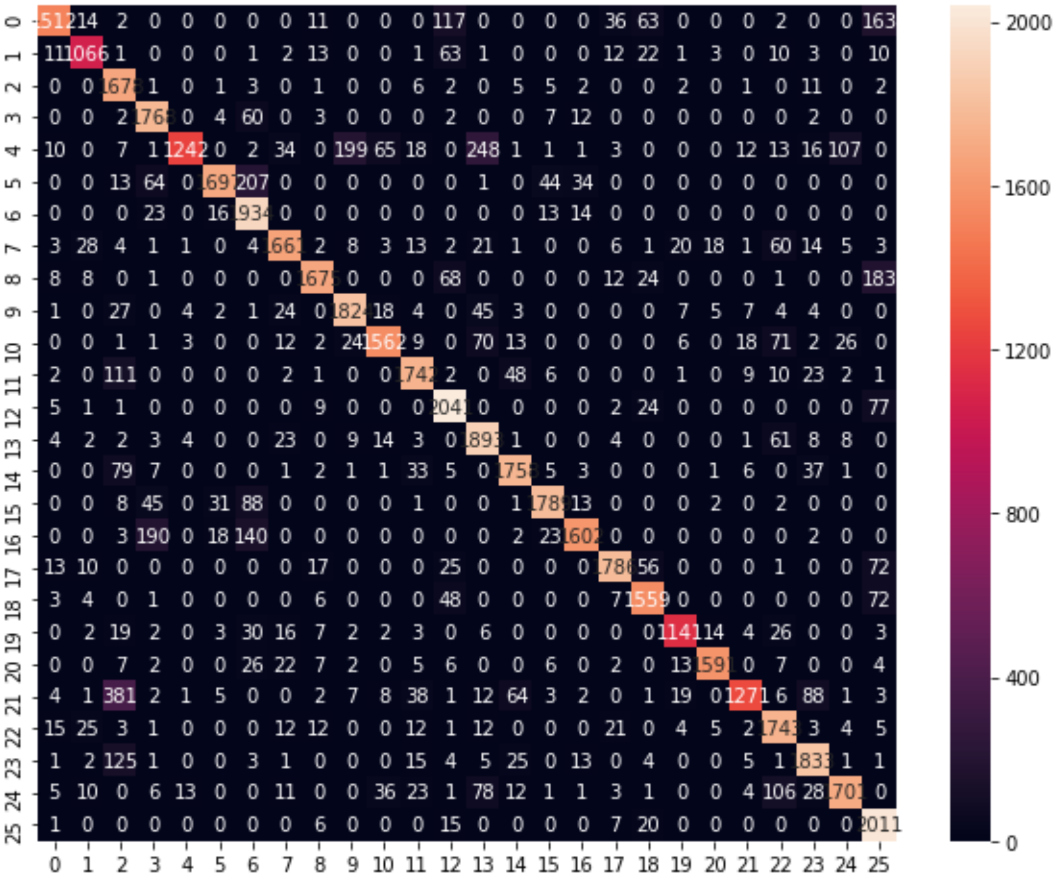
